# Deep learning for fluorescence lifetime predictions enables high-throughput *in vivo* imaging

**DOI:** 10.1101/2025.02.20.639036

**Authors:** Sofia Kapsiani, Nino F. Läubli, Edward N. Ward, Ana Fernandez-Villegas, Bismoy Mazumder, Clemens F. Kaminski, Gabriele S. Kaminski Schierle

## Abstract

Fluorescence lifetime imaging microscopy (FLIM) is a powerful optical tool widely used in biomedical research to study changes in a sample’s microenvironment. However, data collection and interpretation are often challenging, and traditional methods such as exponential fitting and phasor plot analysis require a high number of photons per pixel for reliably measuring the fluorescence lifetime of a fluorophore. To satisfy this requirement, prolonged data acquisition times are needed, which makes FLIM a low-throughput technique with limited capability for *in vivo* applications. Here, we introduce FLIMngo, a deep learning model capable of quantifying FLIM data obtained from photon-starved environments. FLIMngo outperforms other deep learning approaches and phasor plot analyses, yielding accurate fluorescence lifetime predictions from decay curves obtained with fewer than 50 photons per pixel by leveraging both time and spatial information present in raw FLIM data. Thus, FLIMngo reduces FLIM data acquisition times to a few seconds, thereby, lowering phototoxicity related to prolonged light exposure and turning FLIM into a higher throughput tool suitable for analysis of live specimens. Following the characterisation and benchmarking of FLIMngo on simulated data, we highlight its capabilities through applications in live, dynamic samples. Examples include the quantification of disease-related protein aggregates in non-anaesthetised *Caenorhabditis (C*.*) elegans*, which significantly improves the applicability of FLIM by opening avenues to continuously assess *C. elegans* throughout their lifespan. Finally, FLIMngo is open-sourced and can be easily implemented across systems without the need for model retraining.

## Introduction

Fluorescence lifetime imaging microscopy (FLIM) has become an essential technique for studying biological systems at a molecular level across various fields^1^, including cancer research^2–4^, neurodegeneration^5–8^, and plant science^9–11^. FLIM captures not only spatial information but also changes in fluorescence intensity over time, enabling the extraction of fluorescence lifetime data from every image pixel. The fluorescence lifetime is the average duration a fluorophore remains in the excited state before emitting a photon^1^ and is highly sensitive to the fluorophore’s microenvironment. Local variations of different factors, such as temperature, ion concentration, and protein-protein interactions^8,12–14^, etc., all lead to alterations in fluorescence lifetime. As a result, FLIM can provide readouts of molecular and micro-environmental changes within a sample that often remain hidden in imaging techniques where intensity only is recorded^6^.

The data acquisition in FLIM can be performed in both the time and frequency domains, with Time-Correlated Single Photon Counting (TCSPC) playing a predominant role due to its superior time resolution and high signal-to-noise ratio^15–17^. In TCSPC-FLIM, the sample is excited with a pulsed laser, and the time between the laser pulse and the arrival of the first emitted fluorescence photon at the detector is recorded^18^. The process is repeated for many pulses and a histogram of arrival times informs on the most probable fluorescence lifetime of the excited state. However, traditionally the arrival times are extracted using methods such as exponential fitting or phasor plots, which require hundreds to thousands of photons to be accumulated per pixel to ensure reliable quantification of the fluorescence lifetime^15,19,20^. This requirement renders TCSPC-FLIM slow, making it difficult to capture dynamic processes in live biological samples and photobleaching, furthermore, prevents analyses over extended time periods, as often required for drug screening applications^21^.

To address this challenge, deep learning methods have been developed to predict fluorescence lifetimes directly from raw TCSPC data at low photon counts. A variety of architectures have been reported, including multi-layer perception (MLP)^22^, 3D convolutional neural networks (CNNs)^23^, 1D CNNs^24^, MLP-Mixer^25^, extreme learning machine (ELM)^26^, generative adversarial networks (GANs)^27^, and ConvMixer^15^. Many of these approaches^22–27^ make sole use of the time dimension only, *i*.*e*., fluorescence decay curves are analysed without considering the spatial information present in the images. Their advantage is that 1D convolutional operations can be used, which are computationally efficient compared to 2D or 3D convolutions^28^. However, a disadvantage is that contextual information is missed that may be present across neighbouring pixels. The latter is crucial in situations where signal-to-noise ratios are poor. Furthermore, many of the published models cannot be easily adopted by other laboratories as the data used for model training and testing contain a system-specific instrument response function (IRF). As a result, these models require extensive retraining and optimisation on a per-instrument basis, a challenge exacerbated by limitations in code availability which can make the implementation of these models difficult.

To address these problems, we introduce FLIMngo, a generalised network for FLIM analysis which is based on the You Only Look Once (YOLO)^29^ architecture (Figure 1). FLIMngo is trained on simulated FLIM data and effectively utilises both the temporal and spatial information present in raw FLIM images. Our model outperforms other existing machine-learning approaches and phasor plot analysis in predicting fluorescence lifetimes from decay curves with low photon counts (10-100 photons per pixel). In particular, FLIMngo accepts raw TCSPC-FLIM data as input and outputs fluorescence lifetime maps reporting the average fluorescence lifetime of the fluorophores in each pixel. This eliminates the need for pre-existing knowledge of the fluorophore’s decay behaviour, which is a crucial parameter required for the exponential fitting of decay curves. Furthermore, since the training data incorporates a range of different IRFs, FLIMngo is capable of making predictions without requiring an instrument-specific IRF when used to analyse data from different setups, as the model is able to extract system properties directly from the input data themselves. Finally, to demonstrate its potential to transform TCSPC-FLIM into a high-throughput technique, we apply FLIMngo for the analysis of FLIM images of live, dynamic *C. elegans* to study disease-associated protein aggregation.

**Figure 1.**
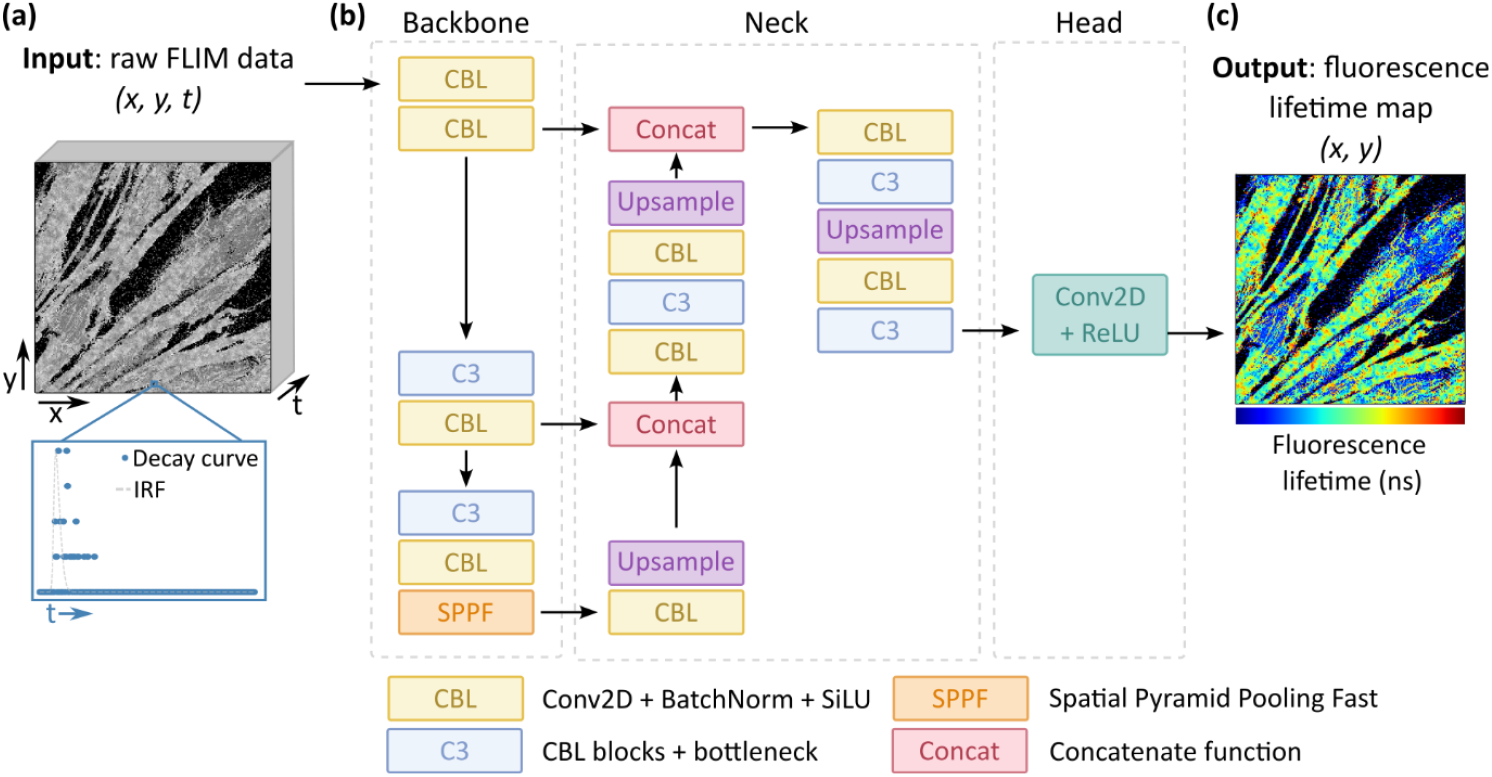
Working principle of FLIMngo. **(a)** The model accepts raw 3D FLIM data as input, where *x* and *y* denote spatial dimensions, and *t* represents time. The time dimension specifically reflects the decay of fluorescence intensity, *i*.*e*., photon counts over time. A low-photon-count decay curve is shown in blue dots, while the IRF is depicted in grey. **(b)** FLIMngo follows an encoder-decoder framework based on YOLOv5. The backbone extracts relevant features while progressively downsampling the input. This is accomplished using CBL modules (2D convolutional operations, batch normalisation, and SiLU activation), C3 blocks (CBL modules with bottleneck residuals), and an SPPF module. The neck further refines these features while performing upsampling and concatenating information from different layers. In the head, the object detection modules in YOLO have been removed and replaced with a 2D convolution operation followed by ReLU activation, a widely used activation function for regression tasks. The YOLO architecture has been adapted from Zhai *et al*. (2022)^41^ in Inkscape. **(c)** The resulting 2D fluorescence lifetime map retains the original spatial resolution and reports the average fluorescence lifetime per pixel.

## Results and Discussion

### Implementation of a YOLO-based architecture for fluorescence lifetime analysis

FLIMngo’s architecture is based on the You Only Look Once (YOLO)^29^ framework, a widely used algorithm for object detection tasks^30^. YOLO offers a balance between speed and accuracy, making it well-suited for real-time data analysis^31^, particularly in applications such as self-driving vehicles, where immediate decision is crucial^32^. Its high computational efficiency also makes it a promising alternative to existing 1D fluorescence lifetime prediction models^24–27^, which likewise offer fast processing. Beyond its strong performance in object detection, YOLO has also been successfully implemented for medical image segmentation^31–33^, demonstrating its versatility across various computer vision tasks. Extending its applicability, we have adapted the YOLOv5 architecture for pixel-wise regression tasks and optimised it for fluorescence lifetime predictions.

FLIMngo employs an encoder-decoder structure with U-Net-like^34^ skip connections, with the complete YOLO network consisting of three main components, *i*.*e*., the backbone, the neck, and the head (Figure ^31^. The backbone is responsible for feature extraction and progressive downsampling of the input image ^35^ using CBL blocks (2D convolutional operations, batch normalisation, and sigmoid linear unit (SiLU)^36^ activation), Concentrated-Comprehensive Convolution (C3) blocks^37^ (CBL layers with bottleneck residuals), and the Spatial Pyramid Pooling Fast (SPPF) module, which captures a broad range of contextual information^38^. The neck further refines and processes the features extracted by the backbone^35^, performing upsampling and concatenation of feature maps from different layers^39^. Finally, the head makes the final prediction.

FLIMngo is an adaptation of the YOLOv5 architecture, where the object detection modules have been removed and replaced with a regression-based output layer using a rectified linear unit (ReLU)^40^ activation function, as shown in Figure 1. While YOLOv5 is traditionally used for 2D object detection tasks, FLIMngo is able to handle 3D TCSPC-FLIM data by encoding the time dimension into the channel dimension. Accordingly, this allows FLIMngo to leverage both temporal and spatial information, in contrast to other published models for predicting fluorescence lifetimes which only focus on temporal information.

Model training was conducted on 1,242 synthetic FLIM images with dimensions of 256 × 256 × 256 (time, x, y). The training dataset was simulated based on single-cell fluorescence intensity images from the Human Protein Atlas (HPA)^42^ database, while ensuring that pixels representing similar cellular structures had fluorescence lifetimes within comparable ranges (see Materials and Methods, Simulated Dataset). Fluorescence lifetimes ranged from 0.1 to 10 ns with up to four exponential decay components, and photon counts per non-background pixel varied between 10 and 2,500.

### FLIMngo reliably predicts fluorescence lifetimes for simulated data from low to high photon counts

FLIMngo’s accuracy in fluorescence lifetime prediction was benchmarked against phasor plot analysis and two published deep learning models, *i*.*e*., FPFLI^15^ and FLI-Net^23^. For benchmarking, a dataset with 40 simulated FLIM images was used with “high photon counts” (100 to 2,500 photons per pixel) alongside the same 40 images simulated with “low photon counts” (10 to 100 photons per pixel), with a representative example shown in Figure 2a. The model performance was assessed by calculating the mean square error (MSE) between the predicted fluorescence lifetime maps and the ground truths.

**Figure 2.**
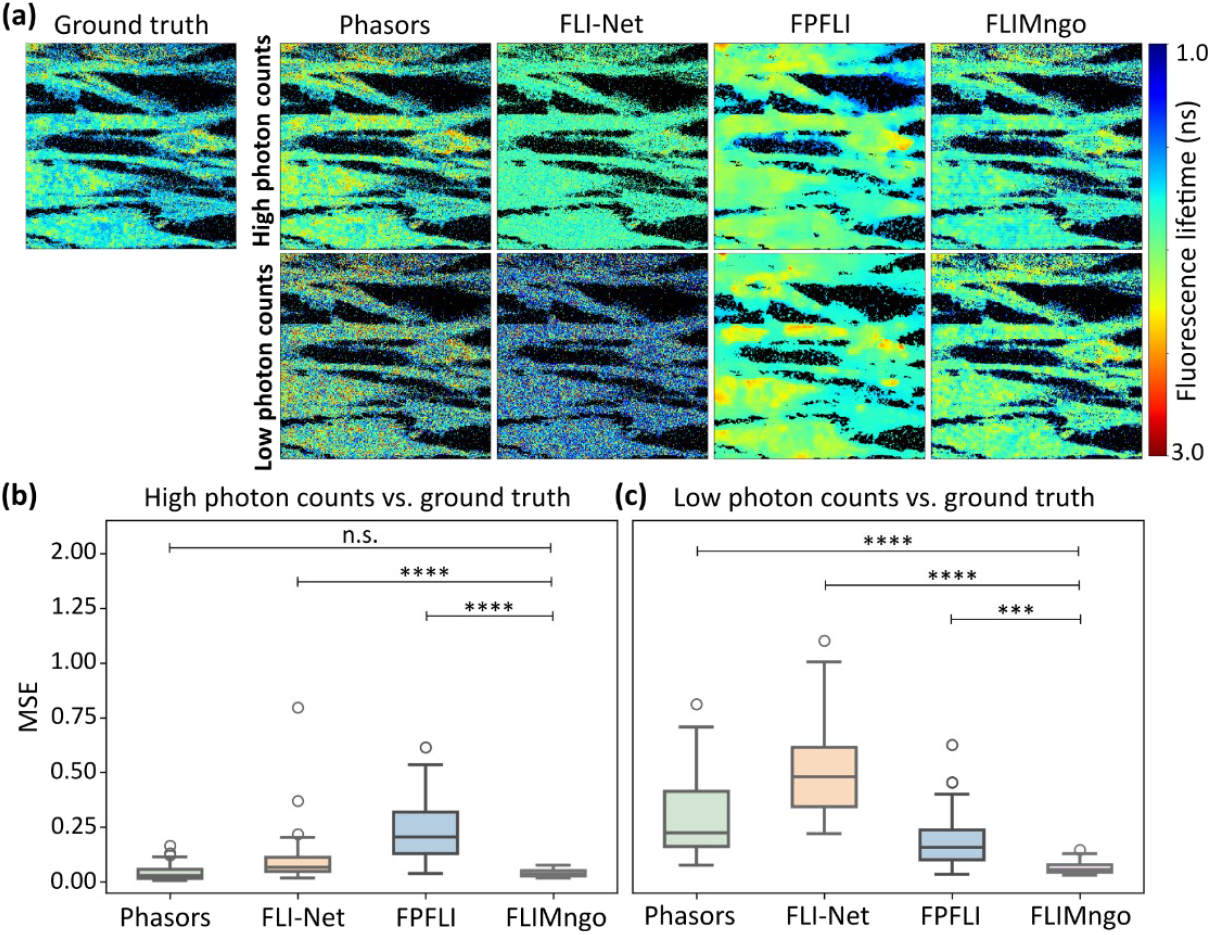
FLIMngo reliably predicts fluorescence lifetimes across photon count conditions. **(a)** Fluorescence lifetime maps of the ground truth alongside the maps predicted by phasor plot analysis, FLI-Net, FPFLI, and FLIMngo. The ground truth fluorescence lifetime maps are identical for both low and high photon counts, as the true fluorescence lifetime value per pixel is independent of photon counts. **(b)** Box-and-whisker plots of the MSE scores for predicted fluorescence lifetime maps from high photon counts compared to ground truth data. **(c)** Box-and-whisker plots of the MSE scores for predicted fluorescence lifetime maps from low photon counts compared to ground truth data. As FPFLI performs pixel binning, the MSE scores were calculated only on pixels representing non-background regions in the ground truths. For the Box-and-whisker plots, the line indicates the median, while the box represents the interquartile range; whiskers extend to the furthest data points within 1.5 times the interquartile range and the dots show outliers^44^. The data consisted of 40 simulated images with high photon counts (100-2500 photons per pixel) and the same images with low photon counts (25-100 photons per pixel), respectively. Statistical significance was calculated using a Kruskal-Wallis test followed by Dunn’s multiple comparisons, where *** denotes p < 0.001, **** denotes p < 0.0001, and “n.s.” denotes non-significant differences.

As presented in Figure 2b, for high photon counts, FLIMngo demonstrated comparable accuracy to phasor plot analysis when quantifying fluorescence lifetimes, thus validating its reliability in photon-rich environments. However, for low photon count data, FLIMngo outperformed all other approaches, as evidenced by the statistically significantly lower mean MSE score between predictions and ground truths (Figure 2c), highlighting its superior accuracy in a photon-starved environment. To further evaluate the effect of binning, we additionally compared FLIMngo to FLIMJ^43^, a decay curve fitting ImageJ plugin, with and without spatial pixel binning, as shown in Supplementary Figure 1. In both cases, FLIMngo had a higher predictive performance for both low and high photon count conditions.

Furthermore, we aimed to establish guidelines on the minimum photon counts required for reliable FLIMngo analysis. Specifically, we simulated 20 FLIM images and repeated the process five times, each with varying photon counts per pixel, *i*.*e*., 100, 80, 50, 20, and 10. Examples of fluorescence decay curves for single pixels can be seen in Figure 3a. As shown in Figure 3b and c, FLIMngo exhibited a much more robust performance across the different photon counts than phasor plot analysis. In particular, FLIMngo maintained a consistent performance down to 50 photons per pixel, whereas predictions became slightly less reliable at 20 photons per pixel and further decreased for 10 photons per pixel (Figure 3d).

**Figure 3.**
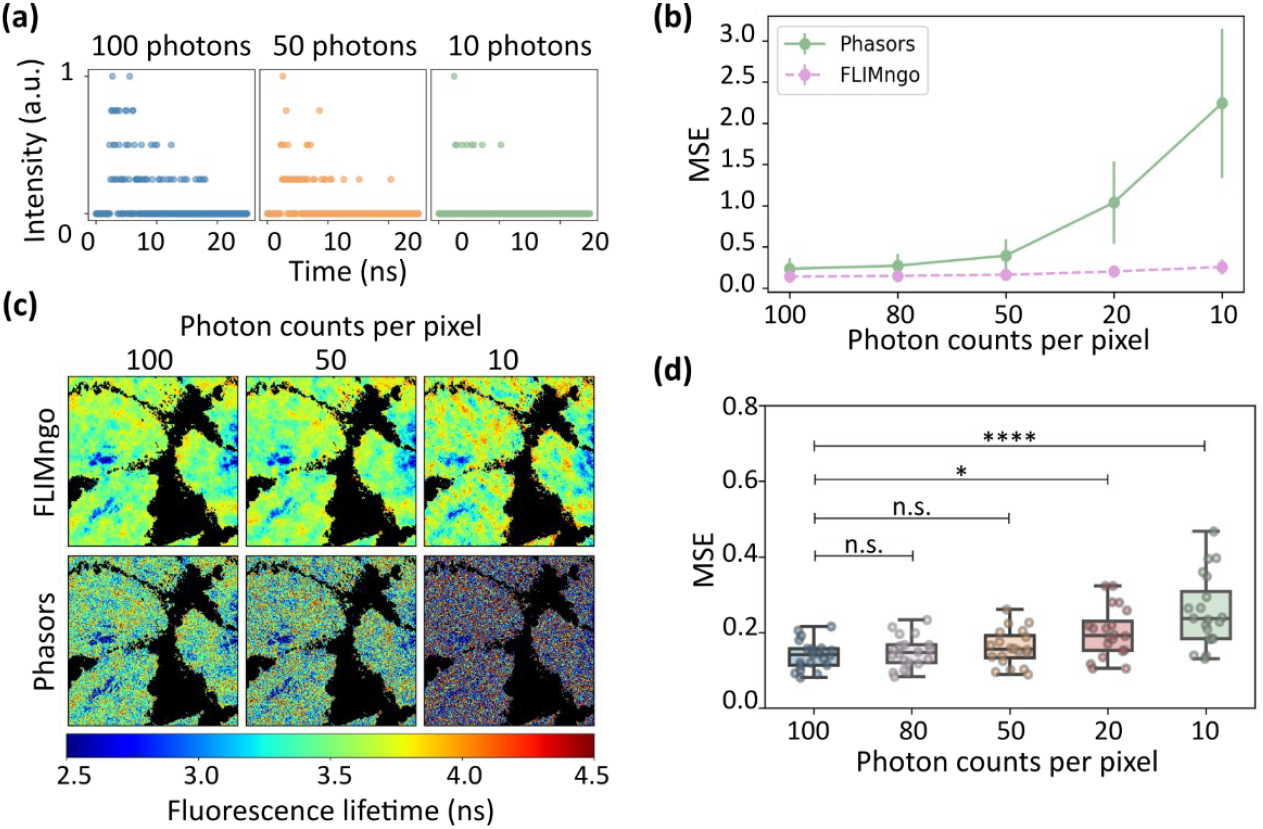
FLIMngo’s performance on simulated low photon count data. **(a)** Example fluorescence intensity decay curves composed of 100, 50, and 10 photons. The decay curves have been normalised to have values between 0 and 1. **(b)** Mean MSE scores calculated between ground truth and predicted fluorescence lifetime maps across varying photon count levels ranging from 100 to 10 photons per pixel. Each photon count condition included 20 simulated FLIM images. The performance of phasor plots is shown in green, while FLIMngo’s performance is shown with a dashed pink line. The error bars indicate standard deviations. **(c)** Fluorescence lifetime maps generated by FLIMngo (top) and phasor plot analysis (bottom) for FLIM images simulated with photon counts of 100, 50, and 10 photons per pixel. **(d)** Box-and-whisker plots of the MSE scores between ground truths and FLIMngo predictions (n = 20). The line indicates the median, while the box represents the interquartile range; whiskers extend to the minimum and maximum values, and individual data points are shown as dots. Statistical significance was calculated using a Kruskal-Wallis test followed by Dunn’s multiple comparisons, with * for p < 0.05, **** for p < 0.0001, and “n.s.” for non-significant difference.

Additionally, to evaluate FLIMngo’s robustness across diverse microscopy setups, we simulated a dataset of 20 high-photon count FLIM images under three distinct IRF conditions. Specifically, each dataset was generated using IRFs with full width at half maximum (FWHM) values of 220 ps, 400 ps, and 800 ps (see Figure 4a) to capture a large range of temporal resolutions. Although TCSPC-FLIM systems typically feature IRFs with FWHMs around 300 ps^45^, a broader IRF was included to assess the model’s performance on data from systems with strongly reduced temporal resolutions or non-ideal configurations.

**Figure 4.**
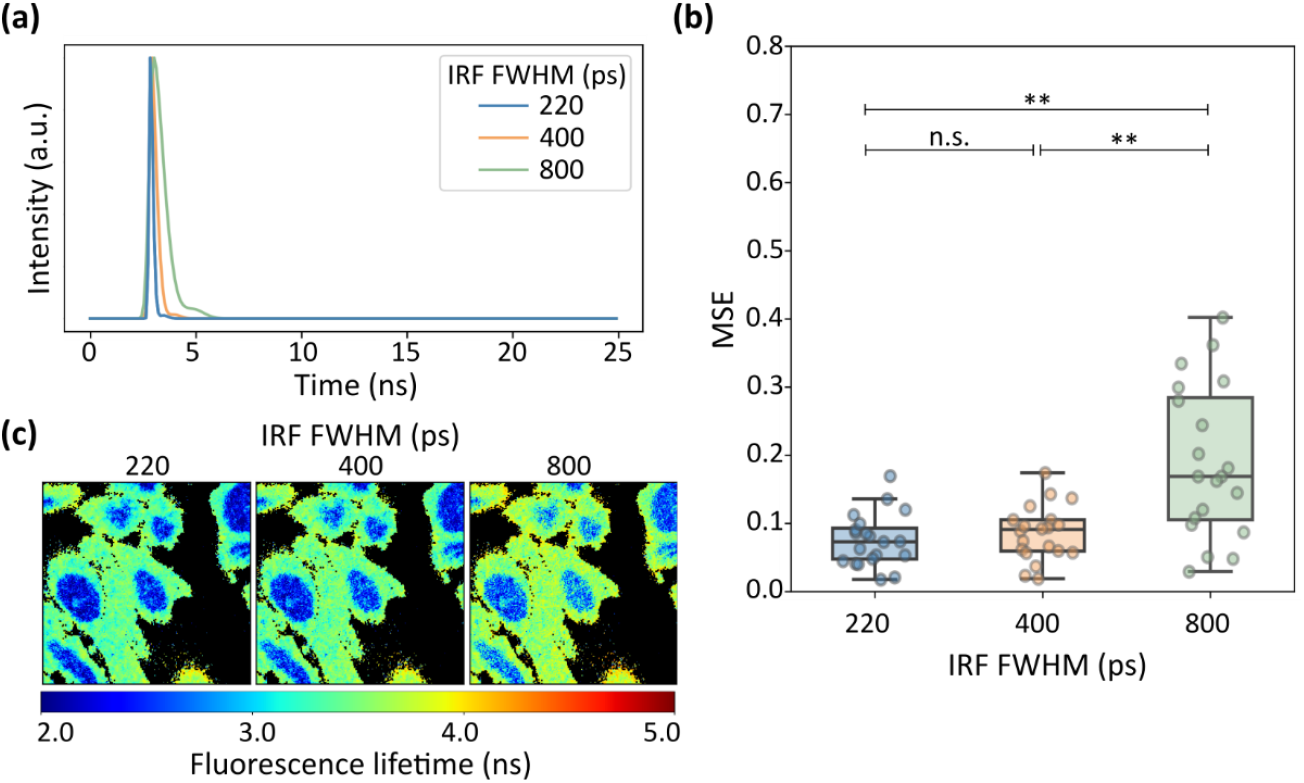
FLIMngo can analyse data simulated with different IRFs. **(a)** Visualisation of IRFs with different FWHM values used in data simulations: 220 ps (blue), 400 ps (orange), and 800 ps (green). **(b)** Box-and-whisker plots of the MSE scores between ground truth and FLIMngo-predicted fluorescence lifetime maps for FLIM data simulated with IRFs of FWHM of 220 ps (left), 400 ps (centre), and 800 ps (right). The horizontal line within each box indicates the median MSE, while the box bounds represent the interquartile range. Whiskers extend to the minimum and maximum values, and individual data points are shown as dots. Each condition included 20 FLIM images simulated with 100 to 2500 photons per pixel. Statistical significance was calculated with a Kruskal-Wallis test followed by Dunn’s multiple comparisons with ** denoting p < 0.01 and “n.s.” denoting non-significant differences. **(c)** Example fluorescence lifetime maps predicted by FLIMngo for data simulated with IRFs of FWHM of 220 ps (left), 400 ps (centre), and 800 ps (right).

No statistically significant differences in the model’s predictions across datasets generated with IRFs of 220 ps and 400 ps FWHM were observed. However, as illustrated in Figure 4b, there is a clear decrease in the prediction accuracy for very wide 800 ps IRFs. This anticipated reduction in prediction performance is likely attributable to the inherent limitations of broader IRFs which blur the early temporal profile of fluorescence decays, thus making predictions more challenging.

Our results demonstrate FLIMngo’s capability to estimate fluorescence lifetimes reliably across a wide range of photon count conditions, maintaining high accuracy even in photon-limited environments down to 50 photons per pixel. Additionally, the model performs well with most standard TCSPC-FLIM setups, however, in setups with strongly reduced temporal resolution that produce IRFs with an FWHM of around 800 ps, users should expect a potential reduction in prediction accuracy.

### FLIMngo makes predictions using both temporal and spatial information

Beyond the improvements in accuracy demonstrated in low photon count environments, the novelty of our approach stems from the integration of both spatial and temporal information for fluorescence lifetime predictions. This is in contrast to other methods that rely solely on the temporal dimension. To validate that the spatial dimensions are also utilised in our model’s predictions, we simulated 20 FLIM images each with dimensions of 256 × 28 × 28 (time, x, y) where all pixels were assigned an identical mono-exponential fluorescence lifetime value that is randomly selected between 0.4 and 6 ns. For each of these images, a second FLIM image was created by setting all pixel values to zero except for a single pixel at position (10,10), as shown in Figure 5a. FLIMngo’s performance in predicting the fluorescence lifetime of the pixel at position (10, 10) was assessed for both the full FLIM image and the single-pixel FLIM image. Specifically, the MSE score between the predicted fluorescence lifetime of the pixel at (10, 10) and its ground truth value was calculated for both conditions. As illustrated in Figure 5b, FLIMngo’s performance was significantly reduced when predicting the fluorescence lifetime of the pixel in the single-pixel image compared to predicting the fluorescence lifetime of the same pixel in the full image with neighbouring pixels present. This confirms that FLIMngo effectively leverages spatial information from adjacent pixels to enhance its fluorescence lifetime predictions.

**Figure 5.**
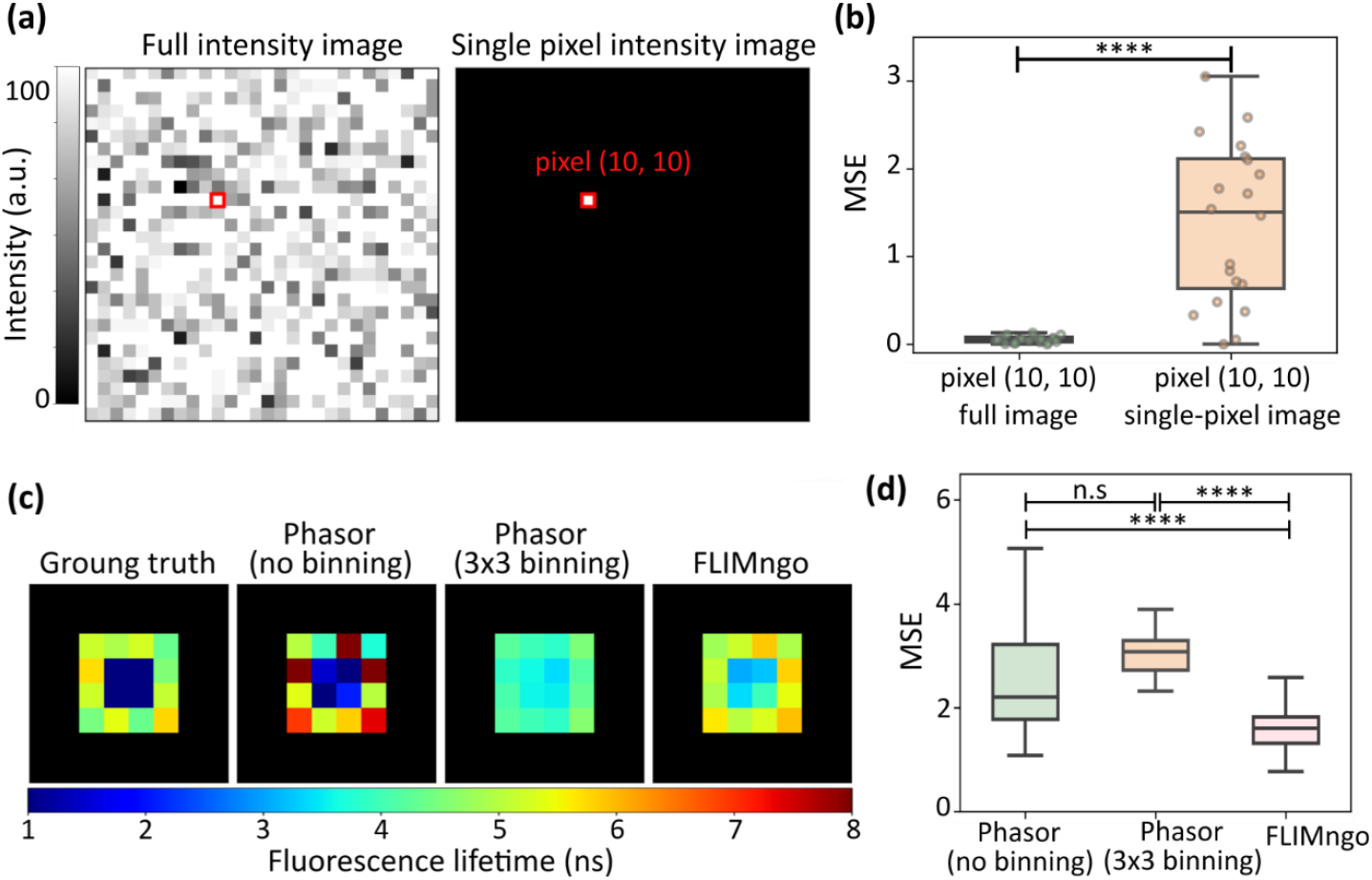
FLIMngo builds on both temporal and spatial information to strengthen its prediction. **(a)** Intensity images for the “full image”, where all pixels are assigned fluorescence lifetimes, and the “single-pixel image”, where pixels are masked to zero except for the pixel at position (10, 10). The pixel at (10, 10) is identical in both cases. The “full image” was simulated with 100 photon counts per pixel, where variations in intensity are due to the addition of Poisson noise. A total of 20 different images were simulated. **(b)** Box-and-whisker plot of the MSE score between the ground truth fluorescence lifetime and the FLIMngo-predicted fluorescence lifetime for the pixel at position (10, 10) in the “full image” (green) and the “single-pixel image” (orange). The line indicates the median, while the box represents the interquartile range; whiskers extend to the minimum and maximum values and individual data points are shown as dots. Statistical significance is calculated using the Wilcoxon Signed-Rank test, where **** indicates a p-value < 0.0001. **(c)** Fluorescence lifetime maps with “flower” structures. **(d)** Box- and-whisker plot of MSE scores between fluorescence lifetime maps predicted for high and reduced photon count data by phasor plots (left) and FLIMngo (right). Statistical significance was calculated using the Mann– Whitney test with **** indicating a p-value < 0.0001.

Additionally, to better understand how FLIMngo strengthens its predictions, we explored whether it incorporates spatial information from adjacent pixels through conventional pixel binning as commonly applied in traditional FLIM analysis methods by comparing its performance to phasor plot analysis with and without 3 × 3 pixel binning. For that, a total of 40 images with 20 photon counts per pixel were simulated. Each image contained a “flower” pattern where the four central pixels had a fluorescence lifetime of 1 ns and the surrounding pixels exhibited lifetimes ranging from 4 ns to 6 ns, as illustrated in the ground truth map in Figure 5c.

FLIMngo outperformed phasor plot analysis both with 3 × 3 and no pixel binning as evidenced by the significantly lower MSE value when compared to the ground truth data (Figure 5d). Specifically, phasor plot analysis with 3 × 3 pixel binning averaged the fluorescence lifetimes across the central and peripheral regions, resulting in a loss of spatial resolution. In contrast, phasor plot analysis without pixel binning preserved the spatial differentiation between the central and surrounding regions, however, several pixels were severely under or over-predicted. FLIMngo demonstrated enhanced accuracy in predicting the fluorescence lifetimes of the surrounding pixels while, nonetheless, predicting higher fluorescence lifetimes for the central pixels than the ground truth, thus proposing that the surrounding area can influence the central pixels undesirably. These findings suggest that FLIMngo effectively integrates spatial information from neighbouring pixels in a more sophisticated manner than traditional binning methods. Future work will focus on further refining the information-sharing process, through additional experiments to better understand how the model utilises information from different pixels, as well as optimising the data simulation process and refining the model architecture.

### FLIMngo accurately predicts fluorescence lifetimes in experimental data

FLIMngo’s ability to accurately predict fluorescence lifetimes from experimental TCSPC-FLIM data was evaluated using 45 recordings of different biological samples imaged on our system. To ensure that the model generalises well across diverse datasets, the images included different cell types (HEK and COS-7 cells), various subcellular structures (nuclei, whole cells, and microtubules), different *C. elegans* nematode strains, as well as the fluorescent dye Rhodamine 6G. Figure 6a shows example images from this dataset, specifically GFP-tagged P525L mutation of fused in sarcoma (FUS) proteins, present inside the head ganglia neurons of *C. elegans* as well as inside human embryonic kidney cells (HEK293T) expressing GFP-tagged wild-type FUS in their nuclei. The experimentally acquired images contained at least 100 photons per pixel (Figure 6b) and are referred to as “high photon count” data. Alongside these, a second dataset with “reduced photon counts” was created from the experimentally acquired images by randomly subsampling the decay curves until each pixel contained 10–100 photon counts, as illustrated in Figure 6c.

**Figure 6.**
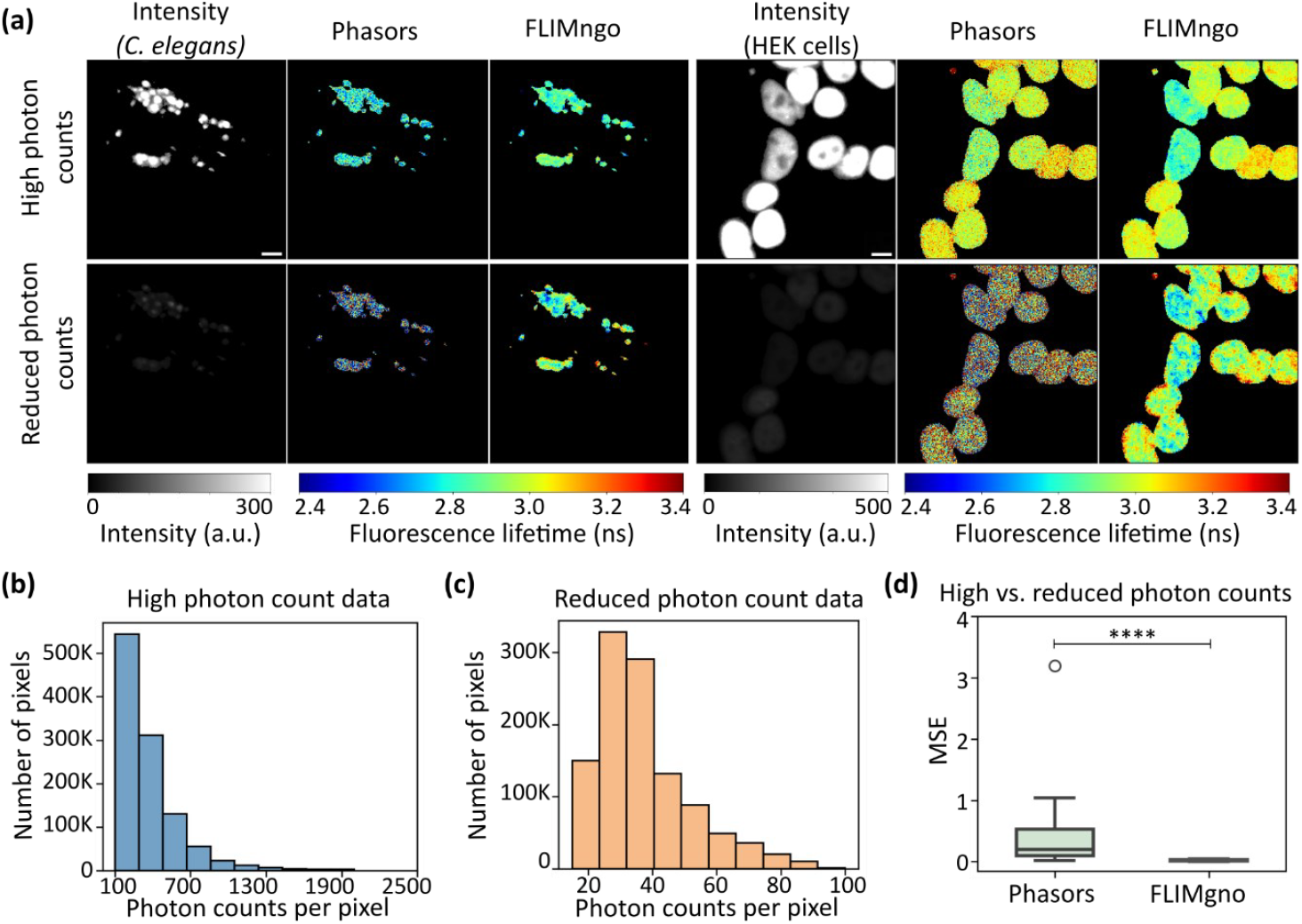
FLIMngo performs well in predicting fluorescence lifetimes in experimental data with low photon counts. **(a)** Example intensity images and predicted fluorescence lifetime maps for experimentally acquired data (“high photon counts”) and the same data with artificially reduced photon counts (“reduced photon counts”). The left side shows *C. elegans* head ganglia neurons expressing the P525L variant of fused in sarcoma (FUS) proteins tagged with GFP. The *C. elegans* have been anaesthetised with levamisole to prevent moving artefacts during imaging. The right side shows HEK293T cells expressing GFP-tagged wild-type FUS in their nuclei. Scale bars are 10 μm. **(b)** Histogram showing the photon count distribution of the high photon count dataset. **(c)** Histogram showing the photon count distribution of the reduced photon count dataset. **(d)** Box-and-whisker plot of MSE scores between fluorescence lifetime maps predicted for high and reduced photon count data by phasor plots (left) and FLIMngo (right). A total of 45 experimentally acquired images were used for the comparison. The line indicates the median, while the box represents the interquartile range; whiskers extend to the furthest data points within 1.5 times the interquartile range and the dots show outliers^44^. Statistical significance was calculated using the Mann–Whitney test with **** indicating a p-value < 0.0001.

To validate that FLIMngo can accurately predict fluorescence lifetimes from experimental data, we compared its predictions on the high photon count dataset to results obtained using phasor plot analysis. The mean MSE score between FLIMngo’s and phasor plot calculations was 0.025±0.022, highlighting the reliability of FLIMngo for analysing experimental data. Next, we compared the MSE scores between the model’s predictions on the high and reduced photon count datasets. FLIMngo demonstrated superior performance, as indicated by a statistically significantly lower MSE for fluorescence lifetime maps derived from high and reduced photon count data compared to phasor plot analysis (Figure 6d).

We further evaluated FLIMngo’s performance on a fixed *Convallaria rhizome* specimen, which is a common plant sample used for validating FLIM setups^46^ due to its complex fluorescence decay behaviour and multiple lifetime components. The same region of the specimen was imaged with varying acquisition times, and FLIMngo demonstrated consistent predictions across images acquired with 2-minute, 24-second, and 2-second acquisition times, as illustrated in Figure 7a and b.

**Figure 7.**
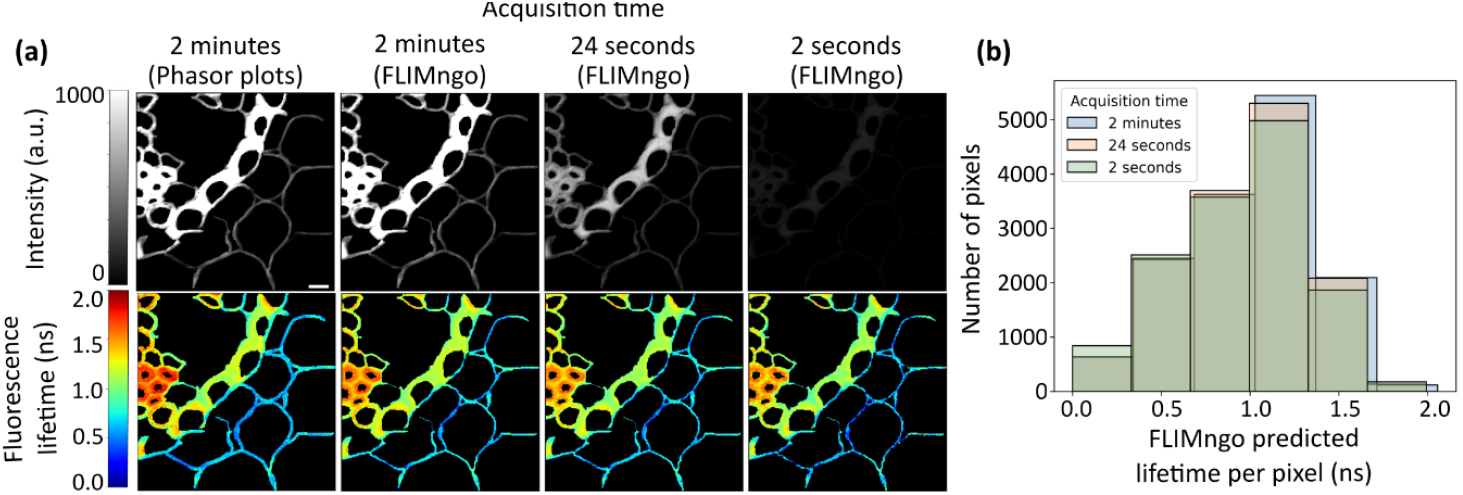
FLIMngo is capable of reliably predicting fluorescence lifetimes in experimental data collected with different acquisition times. **(a)** Intensity images and predicted fluorescence lifetime maps of *Convallaria rhizome* captured with acquisition times of 2 minutes, 24 seconds, and 2 seconds. The 2 minutes recording analysed via phasor plots is shown as a reference. **(b)** Histogram displaying the distribution of FLIMngo predicted fluorescence lifetimes for images acquired with 2-minute (blue), 24-second (orange), and 2-second (green) acquisition times. The scale bar is 10 μm.

Finally, to further evaluate the impact of setup variations, a Rhodamine 6G sample was imaged using two different in-house detectors, namely the photomultiplier tube (PMT) and the avalanche photodiode (APD). The detectors have distinct IRF profiles, with FWHMs of approximately 307 ps and 824 ps, respectively, as shown in Supplementary Figure 2. The mean fluorescence lifetimes predicted by FLIMngo were 3.9 ns for the PMT data and 4.0 ns for the APD data, which fall within the reported fluorescence lifetime range for Rhodamine 6G^47^. This demonstrates that, for simple exponential decays such as the mono-exponential decay expected for Rhodamine 6G, FLIMngo can reliably predict the fluorescence lifetimes even if detectors with very wide IRF profiles are applied.

### FLIMngo permits fluorescence lifetime measurements from fast recordings

Thus far, FLIMngo’s capabilities have been demonstrated on stationary samples. However, a key limitation of traditional TCSPC-FLIM imaging is its poor suitability to capture fast processes or quantify moving samples. For example, FLIM studies using *C. elegans* typically rely on immobilising the nematodes through chemical interventions in order to minimise movement during imaging. However, drug-induced immobilisation with anaesthetics such as levamisole acts by disrupting worm physiology^48^ as well as contributing to stress responses during imaging^49,50^, both of which can severely impact subsequent data interpretation in biomedical research. As demonstrated above, FLIMngo enables the accurate estimation of fluorescence lifetimes from decay curves with minimal photon counts, thereby reducing acquisition times from 2 minutes to a few seconds. This significant reduction in acquisition time accordingly allowed for the FLIM imaging of *C. elegans* mounted on agarose pads without the need for any invasive chemical immobilisation.

We employed FLIM to study the aggregation of amyloid-beta (Aβ) expressed in *C. elegans*, a peptide relevant to neurodegeneration. In particular, accumulation and aggregation of Aβ are a key hallmark of Alzheimer’s disease^51,52^. The disease-causing Aβ1?42 peptide, consisting of 42 amino acids, was expressed in the worms pan-neuronally and substochiometrically labelled with the fluorophore mScarlet. To assess the effects of aggregation on fluorescence lifetime, we compared the fluorescence lifetime of the mScarlet tagged to Aβ_1-42_ peptides to that of mScarlet without Aβ_1-42_ expressed in the neurons of a control strain. FLIM has been utilised in the past for quantifying protein aggregation, including the aggregation of Aβ^52^, as increasing protein aggregation brings the fluorophores into closer proximity, thus increasing self-quenching and reducing the detected fluorescence lifetimes^8^.

Consistent with Gallrein *et al*. (2021)^52^, we found that the fluorescence lifetime of mScarlet-Aβ_1-42_ was significantly reduced compared to that of free mScarlet, indicating that the former is in a more quenched state in these adult nematodes. However, in contrast to earlier experiments, FLIMngo was already able to detect this significant reduction in the fluorescence lifetime maps predicted for 1 second acquisitions using unparalysed worms, as shown in Figure 8. While, for each condition, no significant difference between the mean fluorescence lifetimes recorded in 2 minutes and 1 second were detected, as shown in Supplementary Figure 3. In contrast to FLIMngo’s accurate prediction, phasor plot analysis (Supplementary Figure 4) there was a significant increase in the fluorescence lifetime of the untagged mScarlet in the images acquired over 1 second compared to the 2 minute recordings, thus highlighting the technique’s limitations for quantifying fast or dynamic processes.

**Figure 8.**
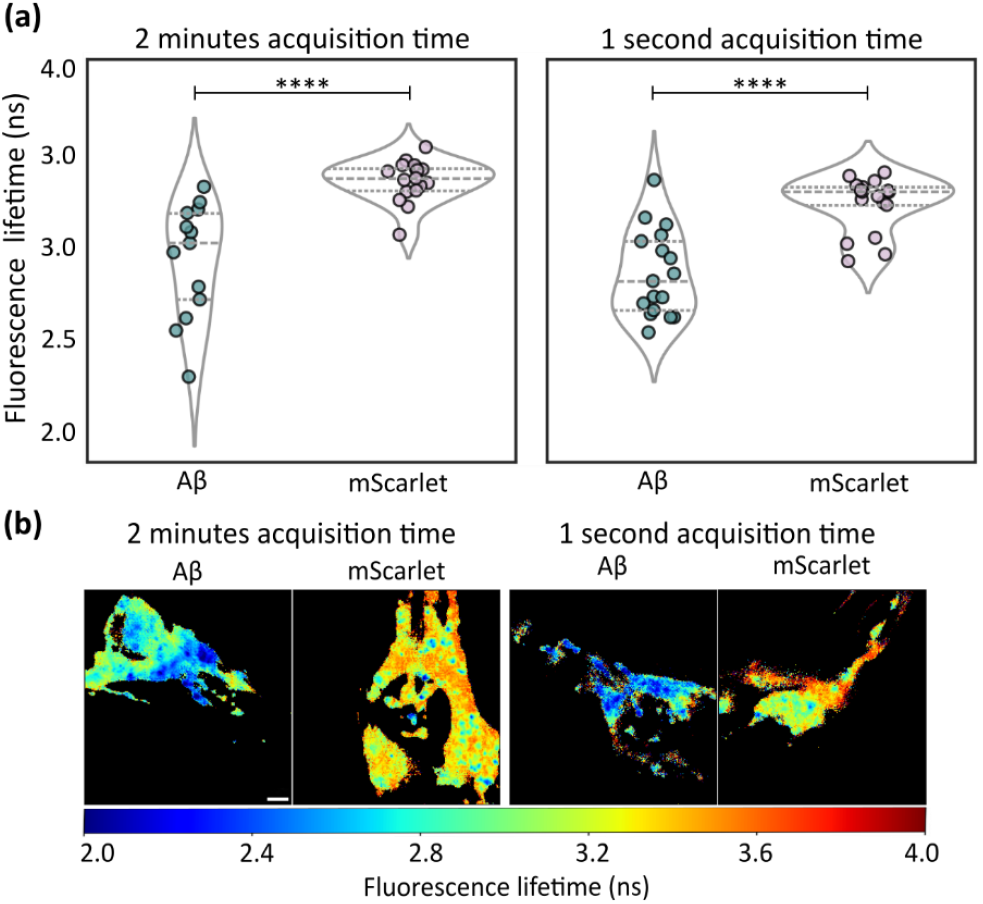
FLIMngo permits the quantification of fluorescence lifetimes in non-anaesthetised *C. elegans*. **(a)** Violin plots comparing FLIMngo-predicted fluorescence lifetimes of mScarlet tagged to Aβ1-42 and mScarlet expressed without Aβ1-42 for data acquired over 2 minutes on paralysed worms (left) and 1 second on freely-moving worms (right). The lines within the plots indicate the interquartile range and median. Statistical significance is calculated using Mann–Whitney tests, where **** indicates a p-value < 0.0001. Images were acquired across two experimental repeats, with a total of 19 images for Aβ1–42 and 20 images for mScarlet under the 1-second acquisition condition, and 19 images for Aβ1–42 and 21 images for mScarlet under the 2-minute acquisition condition. **(b)** FLIMngo-predicted fluorescence lifetime maps for paralysed strains recorded in 2-minute (left) and unparalysed strains with 1-second recordings (right). The scale bar is 10 μm.

## Conclusion

Here, we have introduced FLIMngo, a YOLO-based machine learning model for fluorescence lifetime predictions which leverages both temporal and spatial information present in raw TCSPC-FLIM data, and have demonstrated its capabilities on a diverse set of simulated and experimental data. FLIMngo is open-source and can be applied without the need for model re-training for fluorescence lifetimes ranging from 0.1 to 10 ns and up to four exponential decay components. We have shown that FLIMngo reliably outperforms other approaches, both machine learning-based as well as phasor plot analysis, in quantifying fluorescence lifetimes from low photon count data, *i*.*e*., with 10-100 photons per pixel, while still remaining accurate in high photon count data with more than 100 photons per pixel. Due to its unprecedented performance in photon-starved conditions, FLIMngo is capable of reducing the typical TCSPC-FLIM data acquisition time by more than 100-fold without a loss in prediction accuracy, thus permitting the recording and analysis of dynamic samples. Furthermore, we have provided user guidance to ensure reliable outputs from our model, and have shown that FLIMngo can be successfully employed to quantify data collected across different experimental setups even without the need for an IRF or information on the fluorophores’ exponential decay characteristics. It is further worth highlighting that FLIMngo’s YOLO-based lightweight architecture, similar to alternative 1D models, permits making predictions even without the need for a GPU despite incorporating spatial information.

While FLIMngo can accurately predict data from detectors with IRFs of 100-400 ps FWHM, which is the range found in standard TCSPC-FLIM systems, preliminary results on mono-exponential experimental data show that, with further training and model optimisation, reliable predictions could be made on data collected with detectors of much lower temporal resolutions, such as those with IRFs of 800 ps FWHM, even for highly complex samples. Further, while the current implementation allows for predictions on data with varied spatial dimensions, FLIMngo thus far relies on a fixed time dimension of 256 bins. To account for this, the public GitHub repository provides an option to artificially expand or reduce time dimensions of input data to 256 time bins, thus ensuring FLIMngo remains accessible to a broader range of researchers. Future work will thus involve adapting the model’s architecture to accept data with different temporal dimensions, as well as, further refining the information-sharing process between neighbouring pixels. To summarise, we presented a robust and reliable tool that can significantly reduce the photon count requirements for FLIM data analysis, thereby shortening acquisition times for TCSPC-FLIM measurements and opening new avenues for drug screening and the investigation of dynamic samples and processes.

## Materials and Methods

### Simulated Dataset

Synthetic fluorescence lifetime images with dimensions of 256 × 256 × 256 (time, x, y) were simulated using fluorescence intensity images from the Human Protein Atlas (HPA)^42^ database. The HPA provides single-cell fluorescence intensity images in four colour channels representing microtubules (red), nuclei (blue), endoplasmic reticulum (yellow), and proteins of interest (green). A sliding window was applied to extract 256 × 256 intensity images from the larger HPA images, ensuring that at least 5% of the pixels in each extracted image represented cellular structures, *i*.*e*., non-background pixels. The following process was used to simulate each FLIM image:

#### 1. Assigning Fluorescence Lifetimes

Each colour channel was randomly assigned a base fluorescence lifetime value between 0.1 ns and 10 ns to ensure that similar cellular structures had comparable fluorescence lifetime values.

#### 2. Fluorescence Lifetime Variability

In addition to the base fluorescence lifetime value, each colour channel was assigned a random variability factor ranging from 0.2 ns to 2 ns. This introduced pixel-level variations in fluorescence lifetimes within each channel, which resemble the heterogeneity observed in experimental data.

#### 3. Intensity Scaling

The intensity images from all channels were summed to generate a composite intensity image. This composite image was then scaled to match the target photon count range, 100 to 2,500 photons per pixel for “high photon count” data, and 10 to 100 photons per pixel for “low photon count” data.

#### 4. Simulation of Fluorescence Decay Curves

For each non-background pixel, the fluorescence decay curves were simulated using the following equation *(1)*^24^:

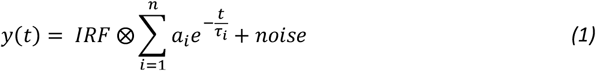

where *IRF* represents the instrument response function and *n* is the number of lifetime components, *i*.*e*., the number of colour channels contributing to the pixel. Additionally, *ai* and *τi* are the fractional contribution and fluorescence lifetime of each colour channel for this pixel, respectively. The term *noise* refers to the Poisson noise typically encountered in TCSPC systems^23^ and ⨂ indicates the convolution between the decay curve and the IRF. The IRF was randomly selected from the collection of simulated and experimental IRFs (see Materials and Methods, IRF Generation). The fractional contribution *ai* of each colour channel was determined using Perlin noise. This technique was used to introduce natural variability^53^ in the contributions of different fluorescence lifetime parameters, as typically observed in experimentally acquired FLIM data. Finally, each fluorescence decay was normalised to have a value between 0 and 1.

Using the approach described above (see Supplementary Figure 5), we successfully simulated FLIM data in which neighbouring pixels exhibit biologically relevant fluorescence lifetimes while also incorporating natural variability as expected to be present in experimental data. The simulated fluorescence decay curves ranged from mono-to quad-exponentials, depending on the contributions from the different colour channels. The ground truth data contained the average fluorescence lifetime for each pixel, and all data were simulated using a bin width of 0.0977 ns and 256 time bins. Nonetheless, FLIMngo is able to handle data acquired at different frequencies by explicitly requiring users to define the bin width (in ns) when importing data. The code for generating the simulated FLIM data is available in FLIMngo’s GitHub repository.

### IRF Generation

To ensure that FLIMngo can accurately predict fluorescence lifetimes across a variety of experimental setups, data simulations were performed using a combination of in-house acquired, publicly available, and simulated IRFs. The in-house IRFs were obtained using both a photomultiplier tube (PMT) and an avalanche photodiode (APD) detector (see Materials and Methods, TCSPC-FLIM setup). Publicly available IRFs were obtained from the GitHub repositories of the FLI-Net^23^, DLTReconstruction^54^, FLIM-fit python library, and FPFLI^15^ projects.

IRFs were simulated to closely match the in-house PMT IRF, as shown in Supplementary Figure 6a. This involved generating a Gaussian-like decay curve^55^ with a slight positive skew to account for the IRF asymmetry due to detector delays. Additionally, an optional secondary peak was introduced to mimic experimental artefacts that are often present in measured IRFs, *e*.*g*., in so-called after-pulsing there can be an additional signal that follows the main detection event a phenomenon commonly observed when single-photon avalanche diode (SPAD)- and PMTs are used as detectors^56,57^. As described by Azzalini (1985)^58^, a skew-normal distribution for a random variable *z* is described by Eq. (2):

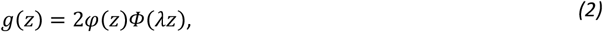

where *z* ∈ (−∞, ∞), *λ* regulates the skewness, *φ*(*z*) is the standard normal density given by Eq. (3):

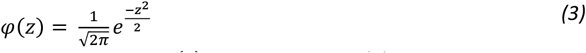

and *Φ*(*λz*) is the cumulative distribution function of *φ*(*z*), described by Eq. (4):

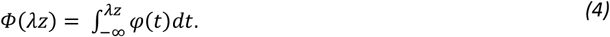

To adjust the peak location and scale of the distribution for the simulated IRFs, we standardised the variable using Eq. (5):

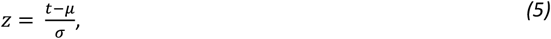

where *t* is the time axis with a bin width of 0.0977 ns and 256 bins, *μ* is the peak location in time, *σ* is the standard deviation calculated as 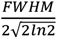. Therefore, the skewed primary peak of the IRF can be described by Eq. (6)^59^, where *u* is a dummy integration variable.:

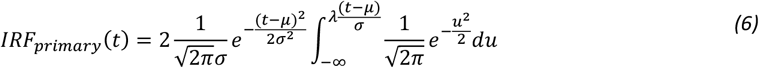

Similarly, the optional secondary peak, simulated to accommodate for after-pulsing, can be modelled by the normal distribution in Eq. (7):

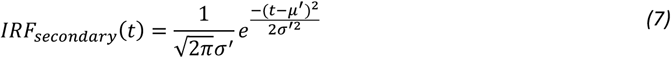

The complete IRF, incorporating both the skewed primary peak and the optional secondary peak, is given by Eq. (8):

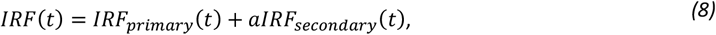

where *a* is the amplitude of the secondary peak, with *a* = 0 removing the secondary peak, signifying that no after pulsing is present. The *IRF*(*t*) was normalised to 1 and IRFs with three different full width at half maximum (FWHM) values, approximately 200, 400, and 800 ps, were simulated both with and without a secondary peak, as shown in Supplementary Figures 6b and 6c, respectively. Finally, to simulate the laser jitter and variations in experimental setup, the IRF peak positions were randomly shifted such that the entire IRF database spans peak positions ranging from the 12^th^ to the 58^th^ time bin (see Supplementary Figure 7).

The bin width of 0.0977 ns was selected as this describes our experimental data when obtained with 40 MHz frequency, however, our model can handle data with different bin widths. The complete set of IRFs, shown in Supplementary Figure 7, exhibited FWHM values ranging from 150 ps to 830 ps. Although IRFs with an FWHM of 800 ps are broader than typically observed, including this extreme case enhanced FLIMngo’s ability to generalise across diverse experimental setups, including non-ideal configurations.

### Model Implementation and Training

FLIMngo was trained on a dataset of 1,242 synthetic FLIM images (see Materials and Methods, Simulated Dataset) split into 80% for training and 20% for validation. The training was performed on an NVIDIA A100-SXM4-80GB GPU, running for approximately 3.5 hours over 150 epochs. The model was optimised using the AdamW optimiser^60^ with a batch size of 10, an initial learning rate of 0.01, and a weight decay of 1e-8. The Mean Squared Error (MSE), was used as the loss function to minimise the error between predicted outputs and ground truth fluorescence lifetime maps (see Eq. *(9)* in Materials and Methods, Model Benchmarking and Evaluation). Early stopping was employed to prevent overfitting, with the training being terminated if the validation MSE did not improve for 20 consecutive epochs. The model was implemented in PyTorch 2.0.1+cu118^61^ and Python 3.11.

### Model Benchmarking and Evaluation

For benchmarking against other models, we retrained FLI-Net^23^ using the Python code available on the project’s GitHub repository. Synthetic bi-exponential FLIM data, based on the MNIST dataset^62^, were generated using our in-house acquired PMT IRFs with fluorescence lifetimes ranging from 0.4 to 4 ns. Two FLI-Net models were trained using data with different photon counts. The first model was trained with 250 and 1500 photon counts and was used for predictions on “high photon count” data, as indicated by FLI-Net’s GitHub repository. The second FLI-Net model was trained with 10 to 100 photons per pixel to assess its performance on “low photon count” data. Moreover, we obtained the pre-trained FPFLI model^15^ from its respective GitHub repository, which had been trained on a fluorescence lifetime range of 1 to 4 ns.

To match FLI-Net’s requirements, models were benchmarked on mono- and bi-exponential simulated data with lifetime ranges of 1 to 4 ns by limiting the colour channels contributing to each pixel to a maximum of two. For the phasor plot analysis, the compared fluorescence lifetime was the average of the modulation and phase fluorescence lifetimes, calculated using FLIMPA^63^. While phasor plot analysis and FLI-Net were assessed on data simulated with an in-house IRF used for FLI-Net training, FPFLI and FLIMngo were evaluated on identical data simulated using an IRF available on FPFLI’s GitHub repository, ensuring comparable conditions. Moreover, as FPFLI performs pixel binning, some of the background pixels were assigned a fluorescence lifetime. Therefore, for this comparison, we only calculated the MSE score between pixels that correspond to non-background regions in the ground truths. Finally, as an alternative analysis to phasors plots, we compared FLIMngo to FLIMJ^43^, an ImageJ plug-in for FLIM data analysis. This enabled the evaluation of FLIMngo against traditional curve-fitting techniques. Specifically, the Levenberg-Marquardt algorithm^64^ was employed, with Poisson as the noise model^65^, two exponential components and our in-house PMT IRF. The analysis was performed both without pixel binning (kernel size = 0) and with 3 × 3 spatial pixel binning (kernel size = 1). In each case, the weighted average of the short and long fluorescence lifetime exponents were reported. Similarly to the FPFLI analysis, due to pixel binning artifacts, FLIMJ’s performance was evaluated only on non-background pixels.

Additionally, FLIMngo’s performance was analysed across varying photon counts (using an PMT IRF measured on our system) and using simulated IRFs with different widths and bi-exponential data simulations with fluorescence lifetimes ranging from 0.4 to 6 ns. Model accuracy was assessed by calculating the Mean Squared Error (MSE) between predicted and ground truth images as defined in Eq. *(9)*^66^:

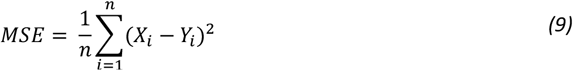

where *m* is the total number of pixels, *Xi* is the predicted value and *Yi* is the truth value for pixel *i*, respectively. An MSE score of zero indicates that the predicted and ground truth images are identical.

### Analysis of Experimental Data

For the phasor plot analysis of experimental data an image of Rhodamine 6G was used as a reference file with a reference fluorescence lifetime of 4 ns. Background intensity masking for experimentally acquired *C. elegans* and human embryonic kidney (HEK293T) cell images was performed by setting the minimum photon counts per pixel to 100 in Python, which is a common photon count threshold used to ensure fluorescence lifetime calculation accuracy^67^. To ensure consistency, identical intensity masks were applied across both high and low photon count measurements.

For the GFP and mScarlet data, fluorescence lifetime maps representing the average phasor fluorescence lifetimes were reported. Intensity masks for *C. elegans* expressing mScarlet were generated for the data collected over 1 second and 2 minutes, by setting a photon count threshold of at least 10 and 100 photons per pixel, respectively. These masks were manually refined using the FLIMfit^68^ software to eliminate regions of autofluorescence and exclude areas containing segments of other worms present in the field of view. A manual mask was created for the *Convallaria Rhizome* (Instruments Direct Services Limited, product code MSAS0321) image using the FLIMfit software, where the same mask was applied to the images taken at different acquisition times.

### Statistical analysis

Statistical analysis was performed in Python (version 3.11.7) using the NumPy (version 1.24.3)^69^, SciPy (version 1.14.0)^70^, Pandas (version 2.2.2)^71^, and scikit-posthocs^72^ libraries (version 0.10.0), while visualisations were created with Matplotlib (version 3.7.1)^44^ and Seaborn (version 0.13.2)^73^.

### Caenorhabditis elegans culture

Normal Growth (NG) plates for *C. elegans* cultures were prepared as described previously^74^. 500 ml of media (for 50 plates) were prepared by dissolving 1.5 g NaCl, 10 g Difco Bacto Agar (Fisher Scientific), and 1.75 g Bacto Peptone (Thermo Fisher Scientific) in 487 ml milli-Q water. Following autoclaving, the solution was kept at 55 °C and 1 M Potassium Phosphate buffer (pH 6.0), 0.5 ml 1 M MgSO_4_, 0.5 ml 1 M CaCl_2_, as well as 0.5 ml 5 mg/ml cholesterol were added whilst stirring. 10 ml of media was added to Petri dishes and allowed to solidify for 24 h. The *Escherichia coli* strain OP50 was cultured in 10 ml nutrient broth in a shaking incubator for 15 h at 37 °C, before 75 µL of OP50 were seeded on each plate.

Experimental images were collected for the following *C. elegans* strains: *C. elegans* pan-neuronally expressing mScarlet (JKM3)^75^ and *C. elegans* pan-neuronally expressing mScarlet tagged to Aβ_1-42_ (JKM5)^75^, both provided by Prof. Janine Kirstein, Leibniz Institute on Aging, Germany. *C. elegans* pan-neuronally expressing GFP-tagged Fused in Sarcoma (FUS-P525L)^76^, provided by Prof. Peter St. George-Hyslop, Columbia University, US.

mScarlet *C. elegans* were synchronised through 6 h egg-laying on day 0 and subsequently transferred to new plates on day 4 (young adults), day 6 (day 3 of adulthood), and day 8 (day 5 of adulthood) to maintain synchronisation. FUS *C. elegans* were synchronised through 6 h egg-laying on day 0 and subsequently transferred to new plates on day 5 (day 2 adulthood) and day 6 (day 3 of adulthood). FLIM imaging of hermaphrodites was performed on day 9 (day 6 of adulthood) for mScarlet strains and on day 7 (day 4 of adulthood) for the FUS strain. For the 1-second recordings of mScarlet control and mScarlet tagged to Aβ_1-42_ *C. elegans* were placed on a 4% agarose pad in a droplet of M9 buffer without anaesthetic treatment, while the 2-minute recordings required *C. elegans* to be immobilised using 5 mM levamisole in milli-Q water. In both cases, *C. elegans* were sandwiched between two coverslips before being mounted onto the microscope for imaging. While this reduces *C. elegans* movement in the z-direction, acquiring high photon count FLIM images is still not possible as the nematodes will still be able to move in the x-y direction.

All strains were kept at 20 °C, while the FUS strain was heat shock treated on day 6 (day 3 of adulthood) and day 7 (day 4 of adulthood) at 33 °C for 2x 30 min each, interrupted by 30 min at 20 °C to induce the phenotype. Treated nematodes were left at 20 °C for at least 30 min following the last heat shock before being used for imaging.

### HEK cell culture

Human embryonic kidney (HEK293T) cells were provided by Prof. Peter St. George-Hyslop^77^ and grown in DMEM (Dulbecco’s Modified Eagle’s Medium, Sigma: D6546) supplemented with 10% heat-inactivated FBS (Fetal Bovine Serum, Life Technology Invitrogen: 10500064), 1% antibiotic-antimycotic mix (Invitrogen: 15240-062), and 2 mM Glutamax-1 (Life Technologies: 35050-038). Cells were engineered with the FUS-expressing gene containing the P525L mutation, GFP-FUS-P525L as described previously^77^. The cell culture was maintained in T75 or T25 cell culture flasks at 37 °C, 5% CO_2_ in air, and 20% humidity. Cells were maintained in the logarithmic growth phase and passaged upon reaching 65-75% confluency (twice per week). For imaging purposes, HEK293T cells were seeded onto Nunc Lab-Tek II chambered cover glass dishes (Thermo Fisher Scientific, 12-565-335) and allowed to grow for 48 hours before being subjected to imaging using FLIM.

### TCSPC-FLIM setup

Imaging was performed on an home built confocal microscope with a TCSPC-FLIM module. Fluorophores were excited using a 40 MHz pulsed supercontinuum laser (Fianium Whitelase, Denmark). The optical setup employed an Olympus IX83 inverted microscope system using a 60x oil objective with a 1.40 numerical aperture (NA) (Olympus, Japan). Photon detection was primarily performed with a PMT (Becker & Hickl PMC-150), and photon counting and data acquisition were managed by an SPC-830 module (Becker & Hickl GmbH, Germany). To assess FLIMngo’s performance on data collected with different detectors, additional measurements were performed using an APD detector (SPCM-AQRH, Excelitas Technologies). IRFs for both PMT and APD were recorded from the back reflection of a blank coverslip. 2-minutes long images were acquired by accumulating photons over 10 cycles with 12 seconds each, 24-second images over 2 cycles, and 1- and 2-second images in single cycles. The laser power was set so that photon count rates did not exceed 1-2% of the repetition rate to prevent photon pile-up.

Filter settings: For GFP-tagged FUS excitation and emission filters were centred at 474 nm and 542 nm (FF01-474/27-25 and FF01-542/27-25, Semrock Inc). Rhodamine 6G, at a concentration of 250 μM in H_2_O, was imaged using filters cantered at 510 nm and 542 nm (FF03-510/20-25 and FF01-542/27-25). For mScarlet, images were acquired using 560 nm excitation and 632 nm centred emission filters (FF01-560/25-25 and FF01-632/148-25) and using a 40x oil objective with a numerical aperture of 1.30 (Olympus). Finally, *Convallaria Rhizome* was imaged using 510 nm and 542 nm centred filters and a 40x oil objective.

## Acknowledgements

G.S.K.S. acknowledges funding from the Wellcome Trust (065807/Z/01/Z) (203249/Z/16/Z), the UK Medical Research Council (MRC) (MR/K02292X/1), Alzheimer Research UK (ARUK) (ARUK-PG013-14), Michael J Fox Foundation (16238 and 022159), and Infinitus China Ltd. C.F.K. acknowledges funding from the UK Engineering and Physical Sciences Research Council (EP/L015889/1 and EP/H018301/1), the Wellcome Trust (3-3249/Z/16/Z and 089703/Z/09/Z), the UK Medical Research Council (MR/K015850/1 and MR/K02292X/1), and Infinitus China Ltd. S.K. acknowledges funding from AstraZeneca and the UK Engineering and Physical Sciences Research Council (EPSRC) grant EP/S023046/1 for the Centre for Doctoral Training in Sensor Technologies for a Healthy and Sustainable Future. S.K. would also like to thank Dr Mona Shehata and Dr Tomasz Witkos from AstraZeneca. N.F.L. acknowledges the Swiss National Science Foundation (Grant Number P2EZP2_199843). We thank Prof. Peter St. George-Hyslop and Prof. Janine Kirstein for providing resources and support for the FUS- and mScarlet-related work, respectively.

## Data availability

All code used to train FLIMngo, along with sample data for model evaluation, is publicly available in the project’s GitHub repository (https://github.com/SofiaKapsiani/FLIMngo).

