## Supplementary Figures for "Deep learning for fluorescence lifetime predictions enables high-throughput *in vivo* imaging"

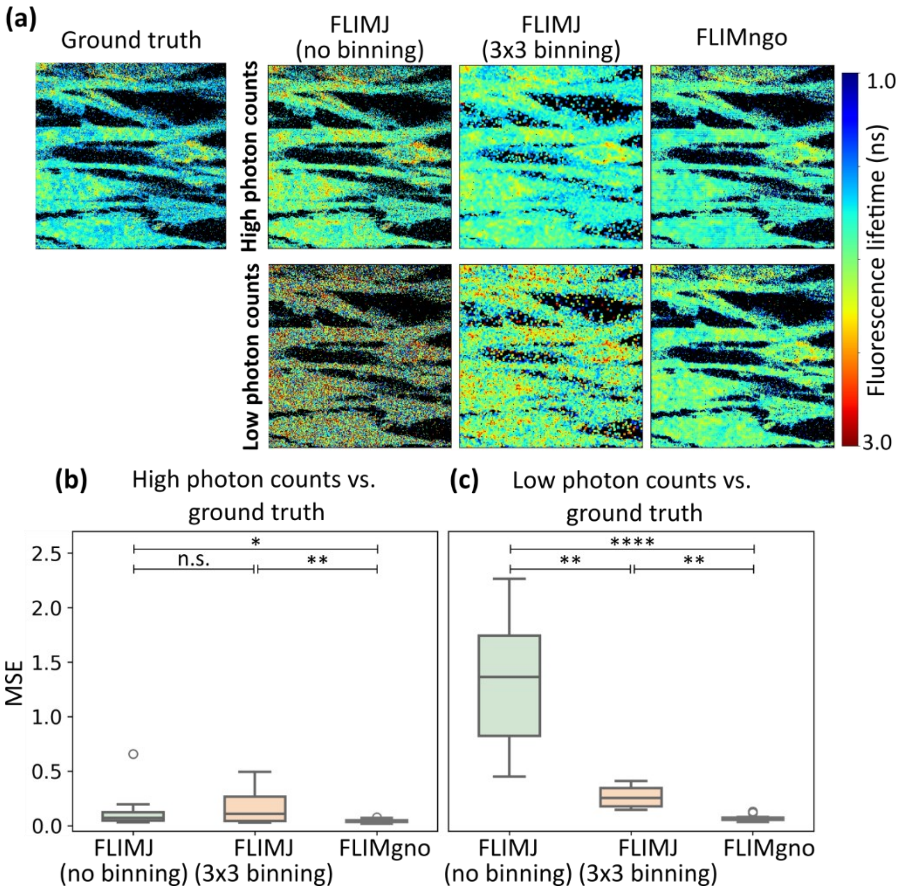

**Supplementary Figure 1. FLIMngo outperforms decay curve fitting on simulated data. (a)** Fluorescence lifetime maps of the ground truth alongside the maps predicted by FLIMJ[1] (no pixel binning), FLIMJ (3x3 spatial pixel binning), and FLIMngo. The high and low photon count datasets have identical ground truth fluorescence lifetime maps. **(b)** Box-and-whisker plots of the MSE scores for predicted fluorescence lifetime maps from high photon counts compared to ground truth data. **(c)** Box-and-whisker plots of the MSE scores for predicted fluorescence lifetime maps from low photon counts compared to ground truth data. For the Box-and-whisker plots the line indicates the median, while the box represents the interquartile range; whiskers extend to the furthest data points within 1.5 times the interquartile range and the dots show outlier. The data consisted of 16 simulated images with high photon counts (100-2500 photons per pixel) and the same images with low photon counts (25-100 photons per pixel), respectively. Statistical significance was calculated using a Kruskal-Wallis test followed by Dunn's multiple comparisons, where * denotes $p < 0.05$, ** denotes $p < 0.01$, **** denotes $p < 0.0001$, and "n.s." denotes non-significant comparisons.

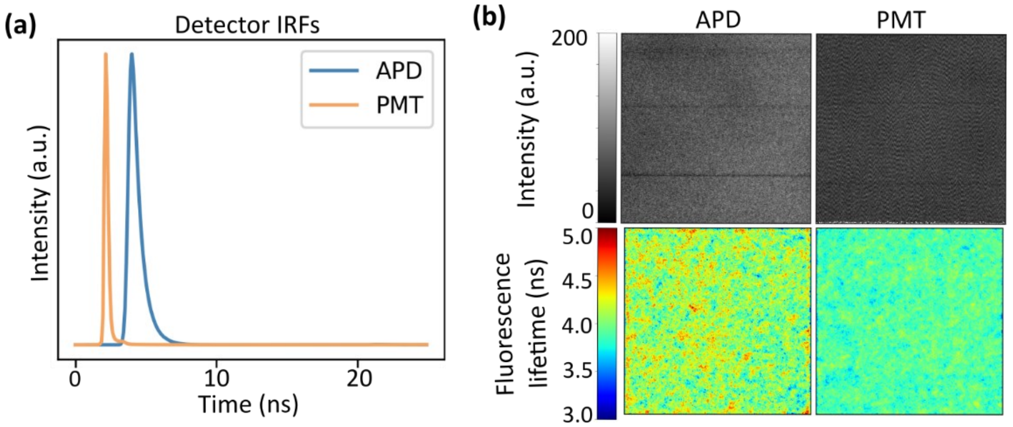

**Supplementary Figure 2. FLIMngo reliably predicts simple exponential decays acquired with detectors having even very wide IRFs. (a)** IRFs obtained using the APD (blue) and PMT (orange) detectors. **(b)** Intensity images and predicted fluorescence lifetime maps for images captured with the APD (left) and PMT (right) detectors.

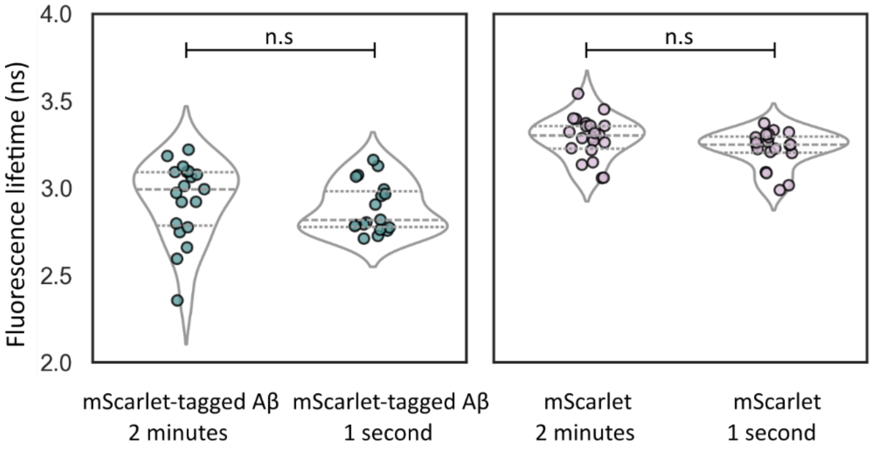

**Supplementary Figure 3. Predicted FLIMngo fluorescence lifetimes for mScarlet expressing *C. elegans*.** Violin plots show FLIMngo predicted fluorescence lifetimes of neuronally expressed mScarlet tagged to $A\beta_{1\text{-}42}$ acquired in 1-second and 2-minute recordings as well as mScarlet without $A\beta_{1\text{-}42}$ imaged in 1-second and 2-minute recordings. The significance levels have been calculated using the Mann–Whitney U test, where “n.s” denotes not significant.

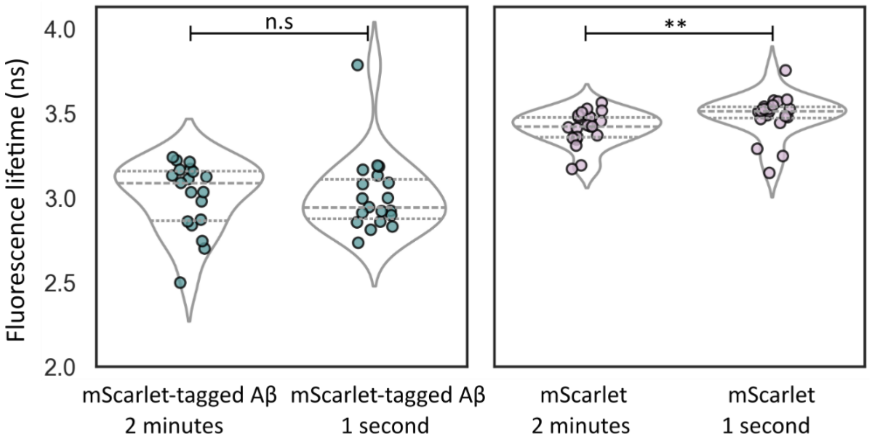

**Supplementary Figure 4. Predicted phasor fluorescence lifetimes for mScarlet expressing *C. elegans*.** Violin plots showing phasor fluorescence lifetimes of neuronally expressed mScarlet tagged to $A\beta_{1\text{-}42}$ acquired in 1-second and 2-minute recordings as well as mScarlet without $A\beta_{1\text{-}42}$ also imaged in 1-second and 2-minute recordings. The significance levels have been calculated using the Mann–Whitney U test, where ** denotes a p-value < 0.0, and "n.s" denotes not significant.

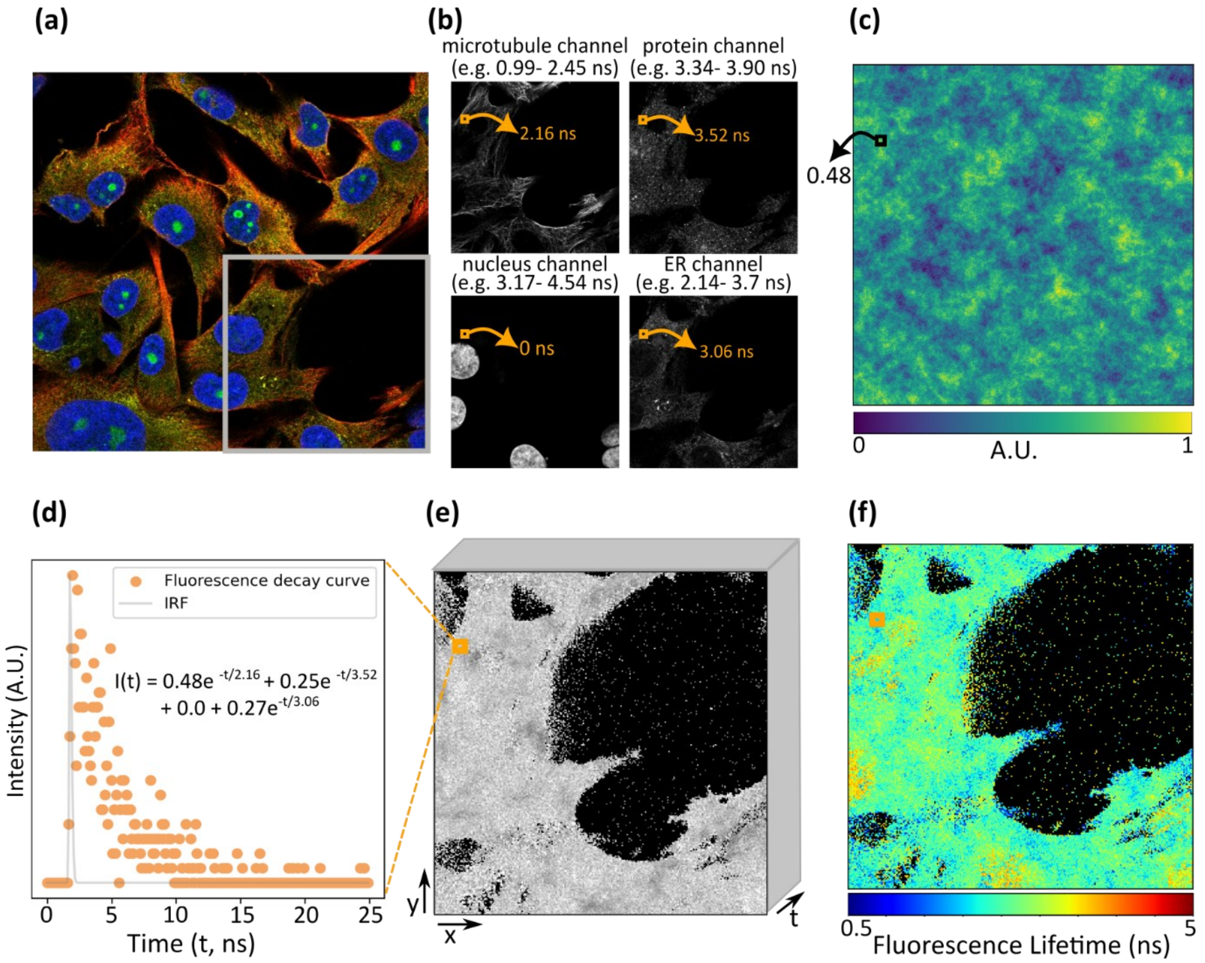

**Supplementary Figure 5. Methodology for simulating TCSPC-FLIM data using the HPA dataset**[2]. **(a)** An example HPA image where the RGBY colour channels represent microtubules, protein, nucleus, and ER, respectively. A sliding window (highlighted in grey) is applied to extract sub-images with 256×256 pixels (x, y). **(b)** Each colour channel is randomly assigned a fluorescence lifetime range, as detailed in Materials and Methods. Example ranges are indicated above each image. For each colour channel, the fluorescence lifetime value for pixel $i$ is displayed in orange. **(c)** Perlin noise is employed to determine the fractional contribution $(a_i)$ of the first colour channel to each pixel. For example, for pixel $i$, $a_i = 0.48$. The contributions of the remaining colour channels to this pixel are then randomly assigned while ensuring that the total fractional contributions sum to 1. **(d)** Example fluorescence decay curve for pixel $i$ generated using $I(t) = I_0 \sum_n a_n e^{-t/\tau_n}$, where $I_0$ is the initial intensity, $a_n$ represents the fractional contribution of the $n$ channel, and $\tau_n$ is its corresponding fluorescence lifetime. The calculation of $I(t)$ from the four channels at pixel $i$ is shown. **(e)** Resulting 3D FLIM image where the *x* and *y* dimensions correspond to the composite intensity image of the four colour channels, while the $t$ dimension contains fluorescence decay curves at each pixel. **(f)** Resulting fluorescence lifetime map where the location of pixel $i$ is highlighted by the orange box.

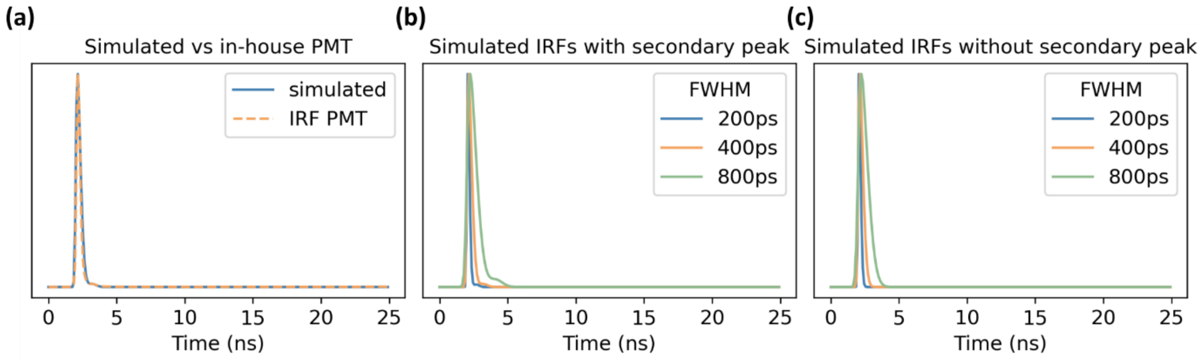

**Supplementary Figure 6. Simulated IRFs and their comparison to in-house data. (a)** Comparison between an experimentally in-house acquired PMT IRF (orange) and a simulated IRF (blue). **(b)** Simulated IRFs with FWHM of approximately 200 ps (blue), 400 ps (orange), and 800 ps (green), with a secondary peak reflecting instrumental artefacts. **(c)** Simulated IRFs with FWHM of approximately 200 ps (blue), 400 ps (orange) and 800 ps (green), without the secondary peak.

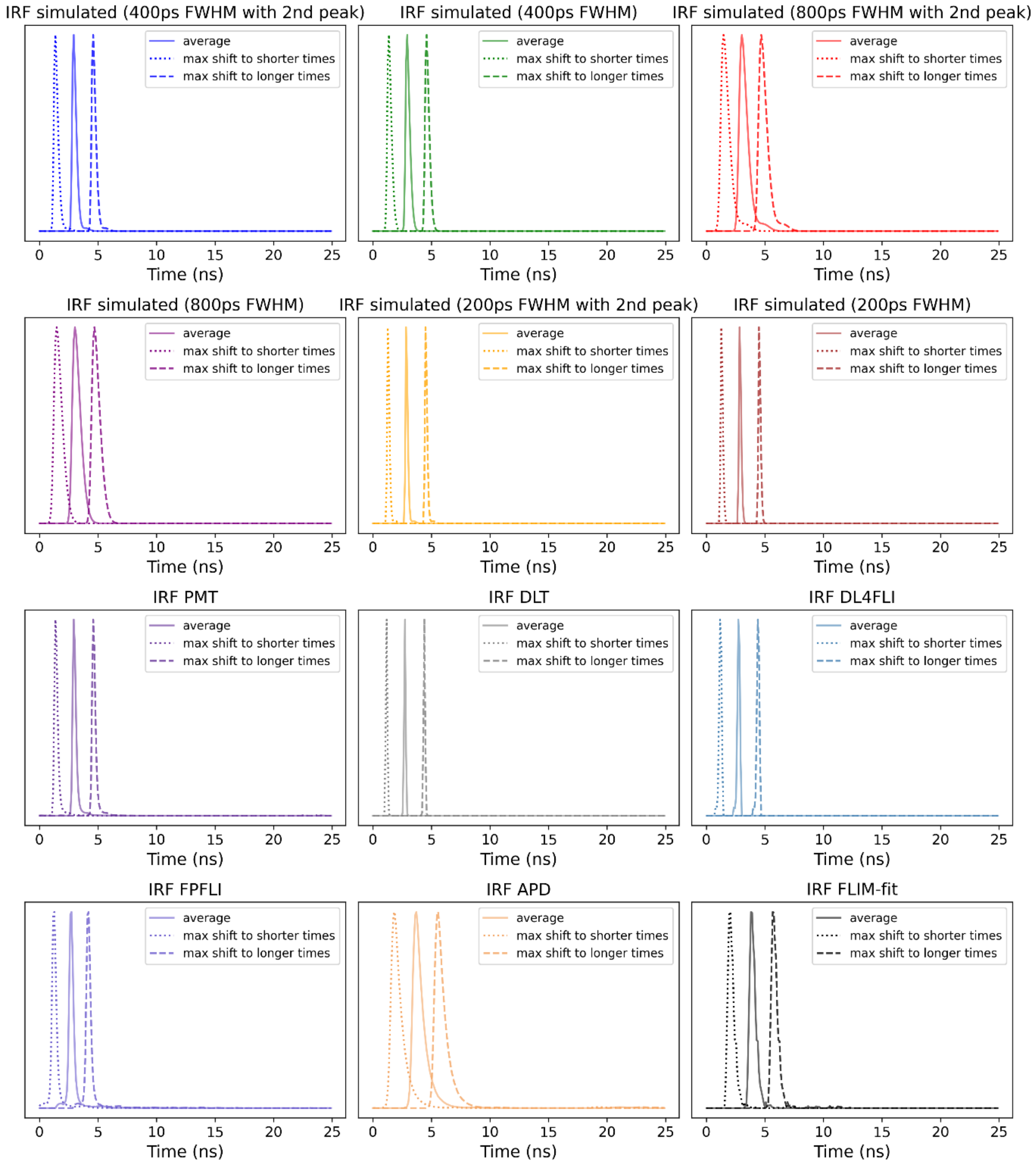

**Supplementary Figure 7. Complete set of IRFs used in the simulation of FLIM data.** To replicate the laser jitter effect often observed in TCSPC-FLIM data, each IRF was randomly shifted to earlier or later time bins. This shifting simulates variability across different experimental setups, enhancing the generalisability of FLIMngo. For each IRF, the maximum shifts towards shorter and longer times are indicated by dotted and dashed lines, respectively, while the average peak position is shown as a solid line. The IRF peak positions spanned from the 12$^{th}$ to the 58$^{th}$ time bin.
